# Unearthing the genes of plant-beneficial marine yeast - *Wickerhamomyces anomalus* strain MSD1

**DOI:** 10.1101/2020.12.22.424010

**Authors:** S Radhesh Krishnan, N Prabhakaran, K Sengali Ragunath, R Srinivasan, K Keerthana Ponni, G Balaji, M Gracy, C Brindha, Lakshmi Narayanan, K Latha

## Abstract

The *de novo* genome of unique marine yeast, *Wickerhamomyces anomalus* isolated from seaweed along Indian coast is presented. The genome assembly was carried out using MaSurCA assembler that generated a data size ∼14.3 mb from short and long reads obtained from Illumina Hiseq 4000 and GridION-X5 respectively. This assembled genome data were used for predicting genes using Augustus gene prediction tool that reported 6720 genes and proteins. The gene sequences were used to unravel the metabolic pathway analysis using KAAS database. The protein sequences were used for secondary analysis to predict the presence of signal peptides using SignalP tool, predicting protein family, domains using Pfam tool and prediction of transmembrane helices in proteins using TMHMM tool. Presence of genes involved in plant growth–promotion and regulation (PGPR) including siderophore and IAA production, iron and sulfur transformation, zinc and phosphate solubilization, nitrogen fixation, synthesis of anti-bacterial and volatile organic compound (VOCs), were assigned. Additionally, acid and alkaline phosphatases, ACC deaminases and lytic enzymes such as β-glucanases, proteases and chitinases involved in pathogen suppression, are also reported. The study elucidates comprehensive understanding of PGP attributes of MSD1 and its potential use in agriculture as bio-fertilizer /bio-stimulant.

## Introduction

*Wickerhamomyces anomalus*, formerly known as *Pichia anomala, Hansenula anomala, Candida pelliculosa* was recently assigned to the *genus Wickerhamomyces* based on phylogenetic analysis of gene sequences, which has caused major changes in the classification of yeasts. This species has been frequently isolated from grapes and wines. *W. anomalus* is a biotechnologically relevant yeast species with food, environmental, industrial, and medical applications [1].

*Wickerhamomyces anomalus* has many different roles in agriculture and the food industry. *W. anomalus* is often among the “film-forming” yeasts associated with beer spoilage [2,3], as well as the spoilage of bakery products [4]. In contrast, *W. anomalus* is among the consortium of yeasts and other microorganisms that are necessary for the fermentation of cocoa and coffee beans, which includes degradation of pectin from the surrounding plant tissue (*Masoud and Jespersen 2006, Schwan and Wheals 2003*). *W. anomalus* has been tested extensively for biocontrol of mold growth that develops during postharvest storage of apples and airtight-storage of grain [5,6]. As summarized by [1], *W. anomalus* can grow under conditions of extreme environmental stress, including anaerobiosis, which makes it strongly competitive with spoilage molds under storage conditions.

The yeast has been reported for its glycosidase [7], volatile organic compound production [1] and antimicrobial [8,9] properties. This species has gained considerable importance for the wine industry since it enhances the flavor of wine [2, 5] and produces bioethanol [10]. The current report ascertains de novo genomic DNA of the isolate confirmed as (NCBI Accession number-MF174856, Safe deposit Accession number-NAIMCC-SD-0004) that was present as an epiphyte on the seaweed Sargassum, Mandapam Beach Park, Tamil Nadu, India. Additionally, a patent has been filed for this yeast and its use in agriculture as a Plant growth promoting yeast (Indian Patent application no. 202041036012). Annotation analysis of the whole genome sequencing (WGS) leads us in prediction and identification of key genes that are responsible for the PGPR activity of the strain MSD1.

## Materials and methods

The marine yeast *W. anomalus* strain MSD1 was isolated from the marine macroalgae (Sargassum sp.,) collected from Mandapam Beach Park, Rameswaram, Tamil Nadu, India [11]. MSD1 was one among the potential seaweed associated microbes (our published research [12]) possessing plant growth promoting microbe like character (Data not disclosed here).

### DNA Isolation, Genome Sequencing, and Assembly & Variation Identification and Genome Diversity Analysis

The yeast was cultured in Zobel Marine Broth (HiMedia, Mumbai, India) for 48 h at 30°C with constant shaking (150 rpm). The high quality DNA from the sample was sequenced at Genotypic Technology Pvt Ltd. India, using Hiseq 4000 (Illumina) and GridION-X5 (Oxford Nanopore Sequencing Technology). The short reads (Illumina) and long reads (Nanopore) data were demultiplexed using bcl2fastq and guppy [13] respectively. Hybrid assembly was performed using Illumina and nanopore reads by MaSurCA Hybrid Assembler [14] with standard parameters. The gene and protein sequence prediction from the assembled genome was performed using Augustus tool [15]. The secondary analysis of the protein was carried out using different protein analytical tools (signalp, tmhmm, PfamScan) [16–18]. The metabolic pathways were predicted using KAAS database.

### Phylogenetic aanalysis

Here we used genome sequence data from 17 publicly available yeast genomes representing 17 known major lineages and 2 non-yeast fungal out groups to generate phylogenetic tree. *Wickerhamomyces anomalus* NRRL Y-366-8 (LWUN00000000.1), *Saccharomyces cerevisiae* (JRIV00000000.1), *Babjeviella inositovora* NRRL Y-12698 (LWKO00000000.1), *Suhomyces tanzawaensis* NRRL Y-17324 (LYME00000000.1), *Metschnikowia bicuspidata var. bicuspidata* NRRL YB-4993 (LXTC00000000.1), *Hyphopichia burtonii* NRRL Y-1933 (LYBQ00000000.1), *Ascoidea rubescens* DSM 1968 (LYBR00000000.1), *Candida arabinofermentans* NRRL YB-2248 (LWUO00000000.1), *Tortispora caseinolytica* NRRL Y-17796 (LSKT00000000.1), *Cyberlindnera jadinii* NRRL Y-1542 (LTAD00000000.1), *Hanseniaspora valbyensis* NRRL Y-1626 (LXPE00000000.1), *Ogataea polymorpha* (AECK00000000.1), *Lipomyces starkeyi* NRRL Y-11557 (LSGR00000000.1), *Nadsonia fulvescens var. elongata* DSM 6958 (LXPB00000000.1), *Pachysolen tannophilus* NRRL Y-2460 (LZCH00000000.1), *Pichia membranifaciens* NRRL Y-2026 (AEHA00000000.1), *Saitoella complicata* NRRL Y-17804 (AEUO00000000.1), *Trichoderma reesei* (AAIL00000000), *Trichoderma harzianum* (JOKZ00000000.1).

### Comparative Genomics

Assembled sequence was compared with the reference sequence to know the gene re-arrangements and genome coverage. We used BRIG to have circular genome representation and Mauve to visualize the synteny between reference genome and assembled genome.

### Nucleotide Sequence Accession Number

This BioProject has been deposited in GenBank under accession number PRJNA556347. The sequences obtained in this project have been deposited in the NCBI Sequence Read Archive under the accession numbers SRR10092046, and SRR9822044. https://www.ncbi.nlm.nih.gov/bioproject/PRJNA556347.

## Results

### General genome characteristics

A total of 6.71 million paired-end reads were generated for the marine yeast from Illumina and 0.37 million reads from Nanopore-GridION respectively. Read statistics are given in **Table 1** and **Table 2**.

**Table 1.**
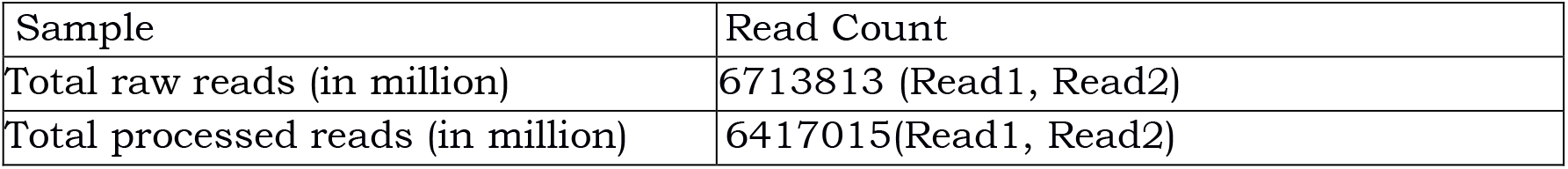
Illumina Read Statistics

**Table 2.**
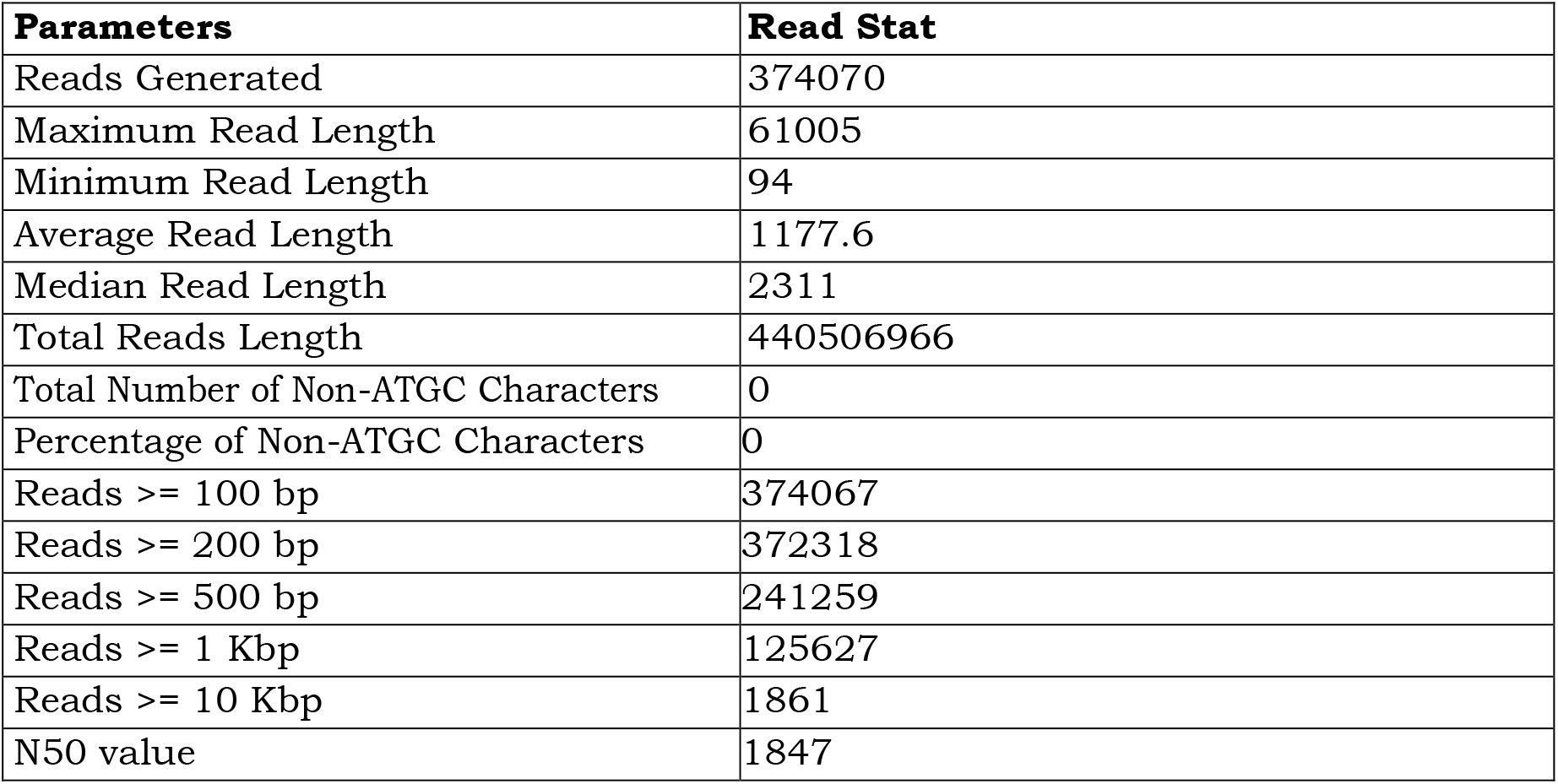
Nanopore Read Statistics

| Parameters | Read Stat |
| --- | --- |
| Reads Generated | 374070 |
| Maximum Read Length | 61005 |
| Minimum Read Length | 94 |
| Average Read Length | 1177.6 |
| Median Read Length | 2311 |
| Total Reads Length | 440506966 |
| Total Number of Non-ATGC Characters | 0 |
| Percentage of Non-ATGC Characters | 0 |
| Reads >= 100 bp | 374067 |
| Reads >= 200 bp | 372318 |
| Reads >= 500 bp | 241259 |
| Reads >= 1 Kbp | 125627 |
| Reads >= 10 Kbp | 1861 |
| N50 value | 1847 |

The size of assembled genome generated was ∼14.3 mb having 289 contigs and the longest contig was of ∼0.2 mb length. Assembly was validated using blast alignment against nr database (**Table 3**).

**Table 3.**
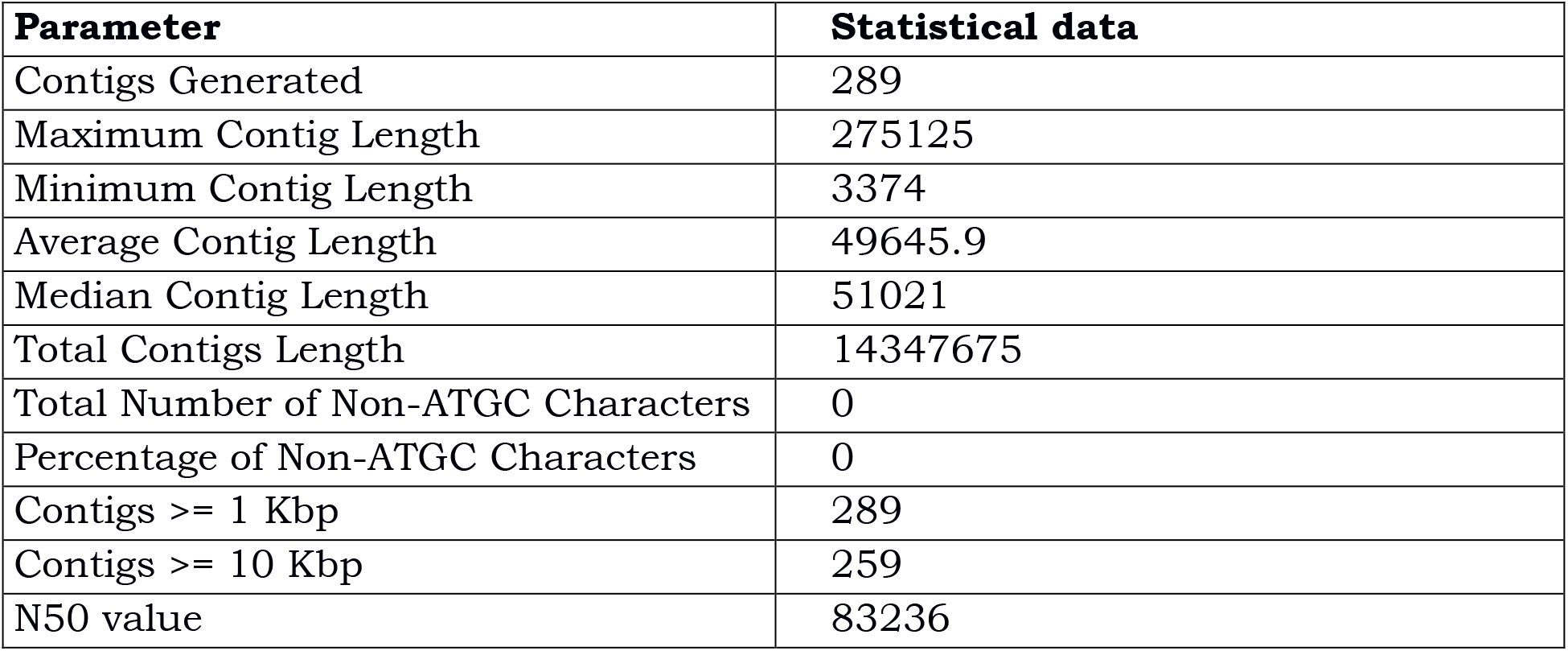
Assembly statistics

The assembled genome sequence was used for predicting genes and protein sequences using Augustus gene prediction tool. A total of 6720 genes and proteins were predicted in the analysis. The GO annotation of the predicted genes was completed using Uniprot database and in-house scripts. Out of 6720 genes predicted, 6658 genes were annotated and 64 genes remained unannotated (**Figure 1**).

**Figure 1.**
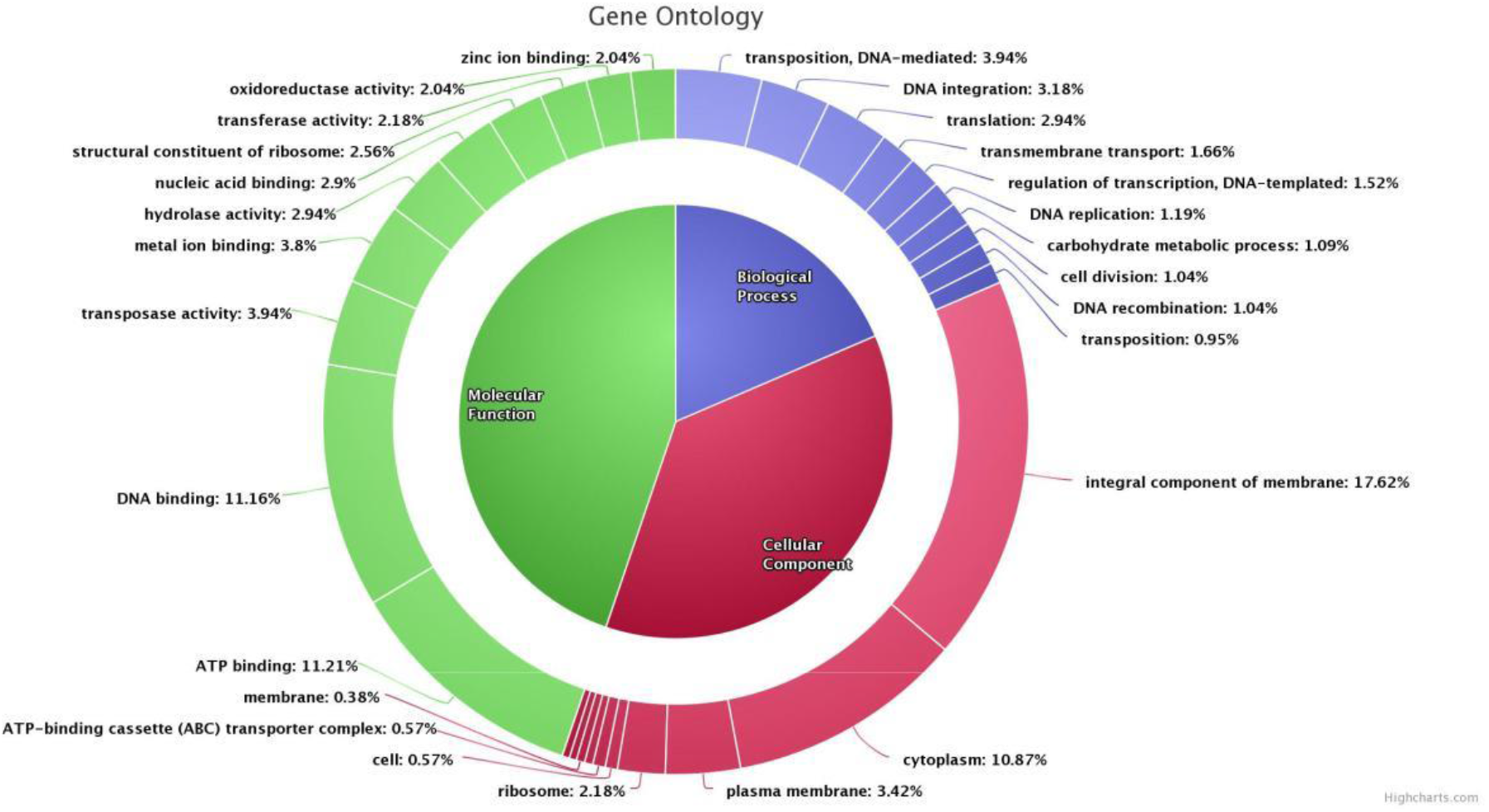
GO Annotation graph

### Comparative Genomics

Reference based whole genome sequencing of *Wickerhamomyces anomalus* was carried out using reference genome available at NCBI for of Wickerhamomyces anomalous, strain-NRRL Y-366-8. More than 140 X of sequencing coverage was achieved for the genome of approximate size 14MB. More than 99% of the reference genome was covered at 1X and >93% of the reference genome was covered at 20 X by good quality data which confirms the choice of reference and the sufficiency of the data for reference based WGS. The consensus sequence resulted from the analysis was compared with the reference sequence in order to know genome significant rearrangements if any (**Figure 2**).

**Figure 2.**
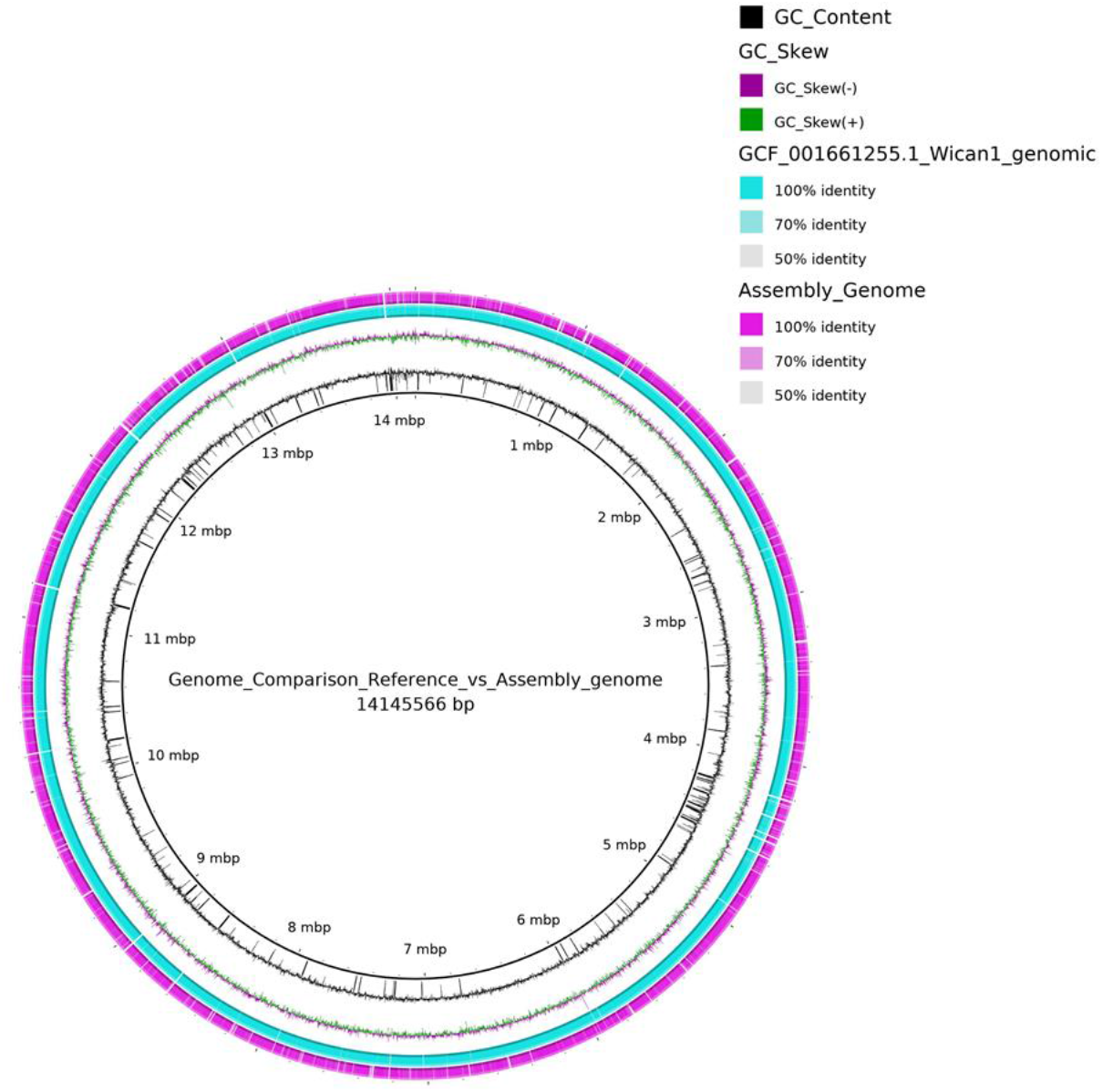
Genome comparison of reference (NRRL Y-366-8) and assembled genome (MSD1)

Assembled sequence was compared with the reference sequence to know the gene re-arrangements and genome coverage. We used BRIG to have circular genome representation and Mauve to visualize the synteny between reference genome and assembled genome (**Figure 3**).

**Figure 3.**
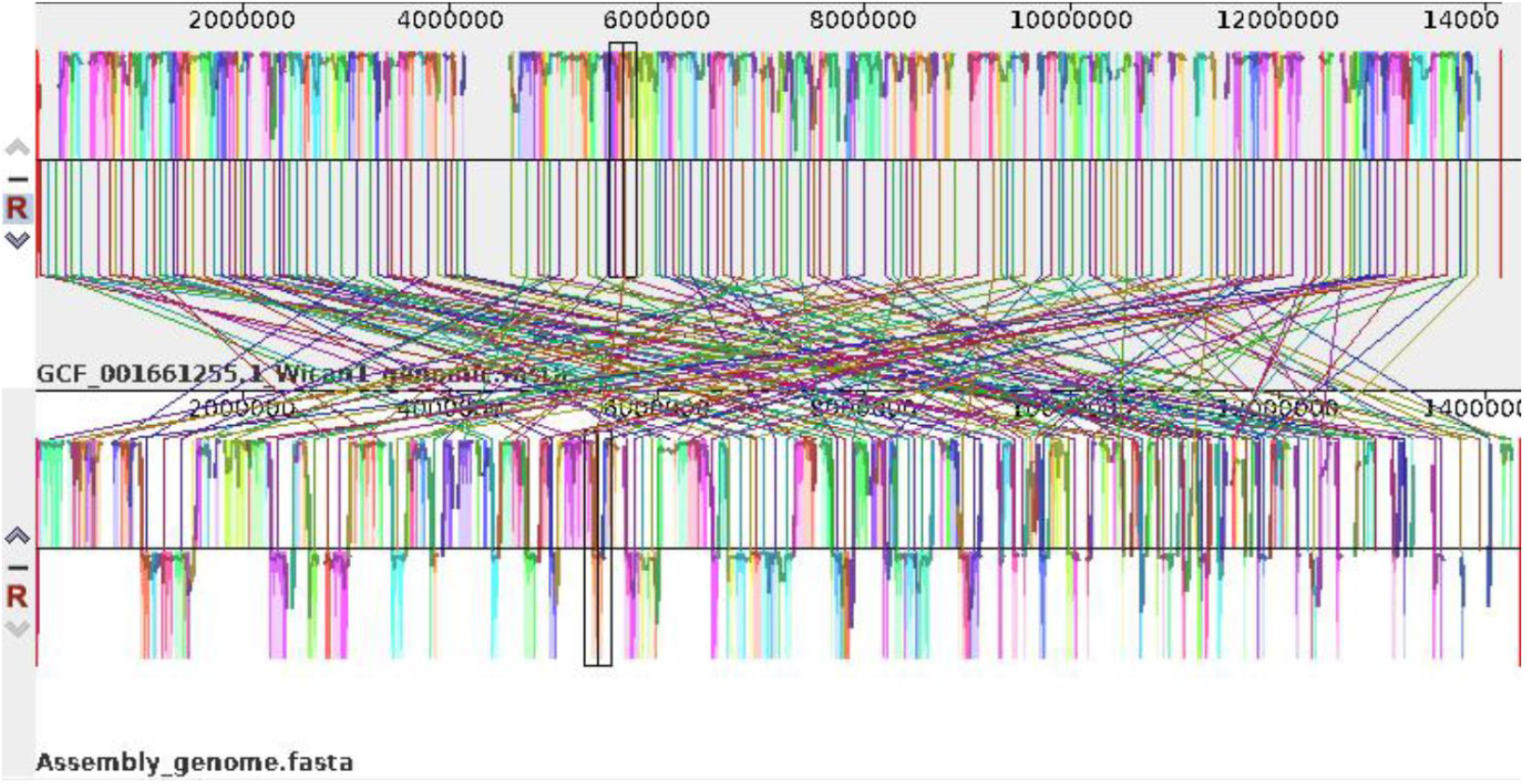
Synteny map of reference (NRRL Y-366-8) and assemble genome (MSD1)

**Figure 4.**
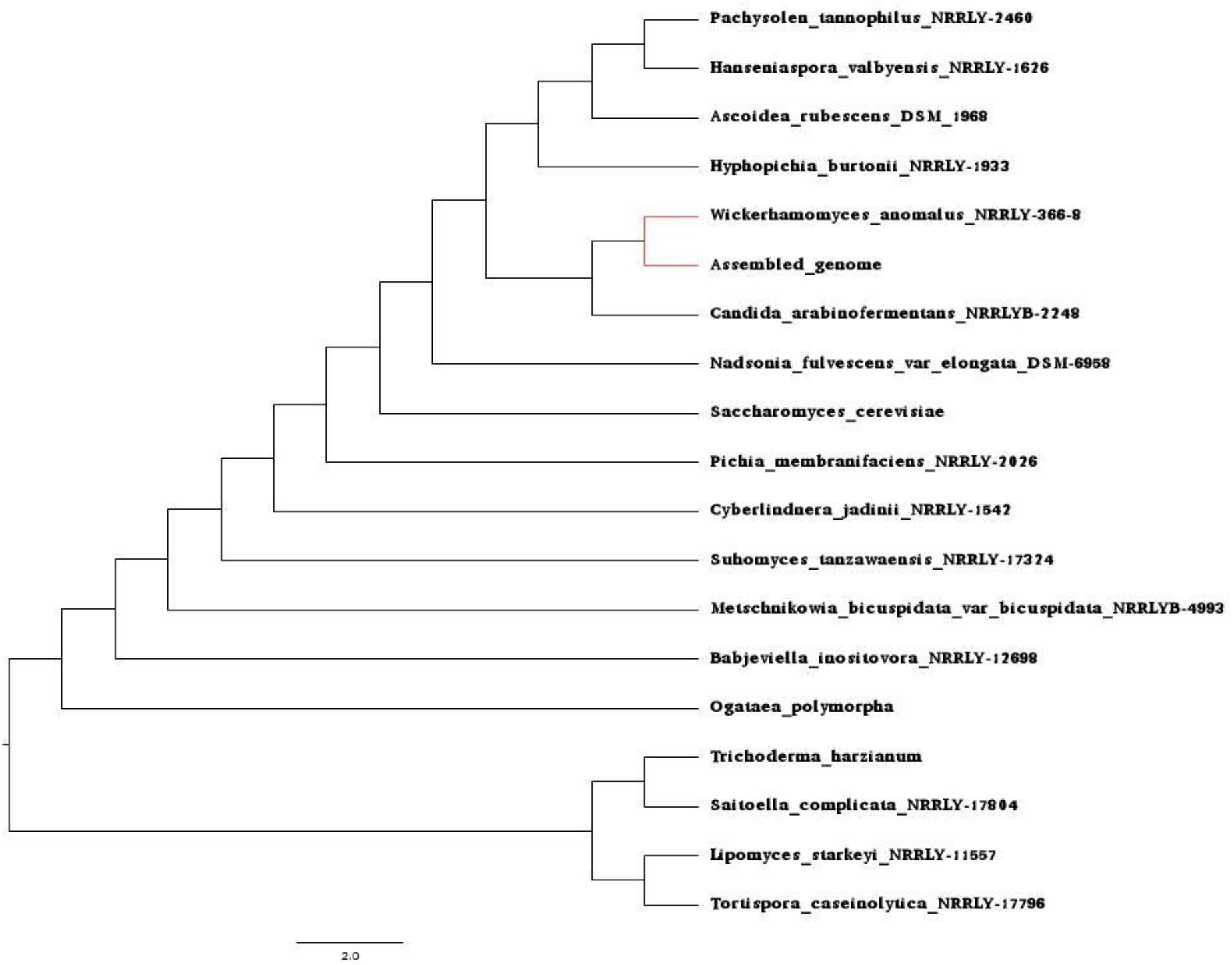
Phylogenetic tree visualizing the comparative genome analysis of MSD1 (Assembled genome) along with other fungal and yeast taxa.

Pathway analysis the yeast genome was carried for using KAAS database that provided functional annotation of genes by BLAST comparisons against the manually curated KEGG GENES database [19]. The result contains KO (KEGG Orthology) assignments and automatically generated KEGG pathways (**Supplementary file 1**). The secondary analysis of protein sequences obtained from Augustus was carried out using different tools-SignalP, Pfam-Scan, TMHMM. SignalP tool predicts the presence of signal peptides and the location of their cleavage sites in proteins. A total of 521 signal peptides were predicted out of which 304 had trans-membrane segments and 217 without trans-membrane segments. Pfam-Scan tool was used to predict protein family and domains present in the predicted protein sequences. A total of 8280 pfam annotation (which includes family, domain, repeat and motif) were predicted for 6720 proteins. Out of 8280 pfam annotation 5339 contained clan (group of related protein families) information and 2941 had no clan information. TMHMM tool predicts transmembrane helices in given proteins sequences. A total of 6720 proteins used for transmembrane helix prediction, out of which 1377 proteins contained transmembrane helices and remaining 5343 proteins were without transmembrane helices (**Supplementary file 2-4**).

## Discussion

### Genes identified from WGS of MSD1 related to PGPR traits

We identified genes in the MSD1 genome attributable to the production of IAA, solubilization of minerals like phosphate and zinc, synthesis of sideropheres, acetoin and 2,3-butanediol, suppression of pathogenic fungi, resistance to oxidative stress, and ability to break down toxic compounds and other abiotic stresses.

Here, two proposed **IAA biosynthesis** pathways, amidase and aldehyde dehydrogenase pathways are identified in the genome of MSD1. In the indole-3-acetonitrile (IAN) pathway IAN can first be converted to indole-3-acetamide (IAM) by nitrile hydratase and then IAM is converted to IAA by amidase. In the IPyA pathway indole-3-pyruvate (IPyA) is converted to indole-3-acetaldehyde (IAAld) by indolepyruvate decarboxylase and then to IAA by aldehyde dehydrogenase. All of these genes responsible for IAA synthesis were present in MSD1 genome [20].

Gluconic acid (GA) is recognized as one of the major organic acids in most bacteria responsible for the solubilization of mineral phosphates. The synthesis of GA is catalyzed by glucose dehydrogenase (GDH) and its co-factor pyrrolo-quinolone quinine (PQQ)[21–23]. Accordingly, the MSD1 genome was searched for the presence for phosphate transporter genes. Gene IDs PHO84 and PHO87 that encodes for inorganic phosphate transport were predicted. In addition, 5 genes encoding mitochondrial thiamine pyrophosphate transporters (solute carrier proteins) were also predicted in the MSD1 genome.

MSD1 carrying the gene encoding for the synthesis of **siderophore** was identified. Genes encoding isochorismate domain containing protein, Gene IDs K08197 (7 copies) and K23503 (2 copies) that are annotated for siderophore-iron: H+ symporter and sideroflexin respectively were predicted. These indicate that although strain MSD1 cannot synthesis numerous sideropheres, it can heterologously obtain siderophores produced by other soil bacteria [20,24].

The MSD1 genome was predicted for the presence of a cascade of genes for Fe uptake/transport. Genes like K07243 (high-affinity iron transporter), K19791 (iron transport multicopper oxidase), K12346 (metal iron transporter), K22736 (vacuolar iron transporter family protein), K15113 (solute carrier family 25), K02304 (precorrin-2 dehydrogenase/sirohydrochlorin ferrochelatase), and K01772 (protoporphyrin/coproporphyrin ferrochelatase) were annotated from MSD1 genome.

### The genes Associated with Plant Growth Promotion Traits

Previous phenotypical and PGP abilities, observed in pure culture and in plant experiments under salt stress, was supported by the MSD1 genome content (**Table 4**). The MSD1-detailed genomic profile of their confirmed PGP abilities and other possible mechanisms involved in plant promotion were analyzed and described here. It has been reported that PGPR may produce compounds such as **phenazine** and 4-hydroxybenzoate which function as antibiotics and suppress plant pathogenic microbes [24]. UbiD, involved in 4-**hydroxybenzoate** synthesis, and PhzC-PhzF, involved in phenazine synthesis, were identified in the MSD1 genome. Moreover, a homologue of the gene coding for **chitinase** enzyme was identified that can potentially dissolve the cell wall of pathogenic fungal and insect pests [24]. In addition to these, the genes gabD and gabT which are responsible for the production of pest/disease inhibiting **γ -aminobutyric acid (GABA)** in the genome was identified [20]. This suggests that the synthesis of the three antimicrobial compounds is a widespread pathway in MSD1.

**Table 4.**
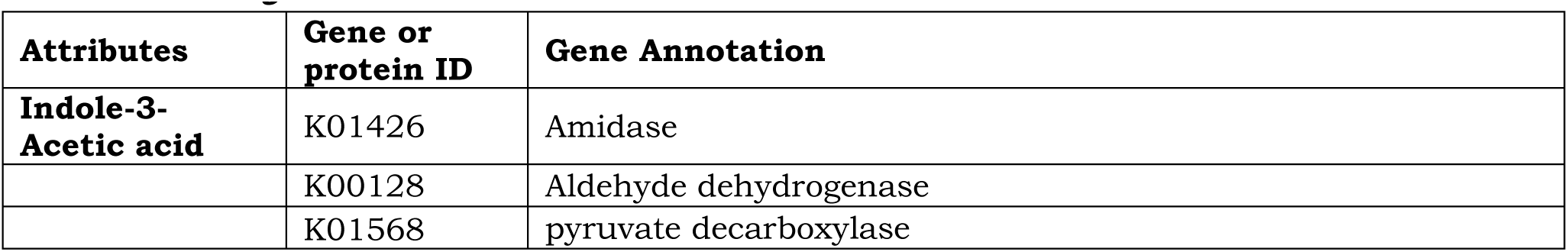
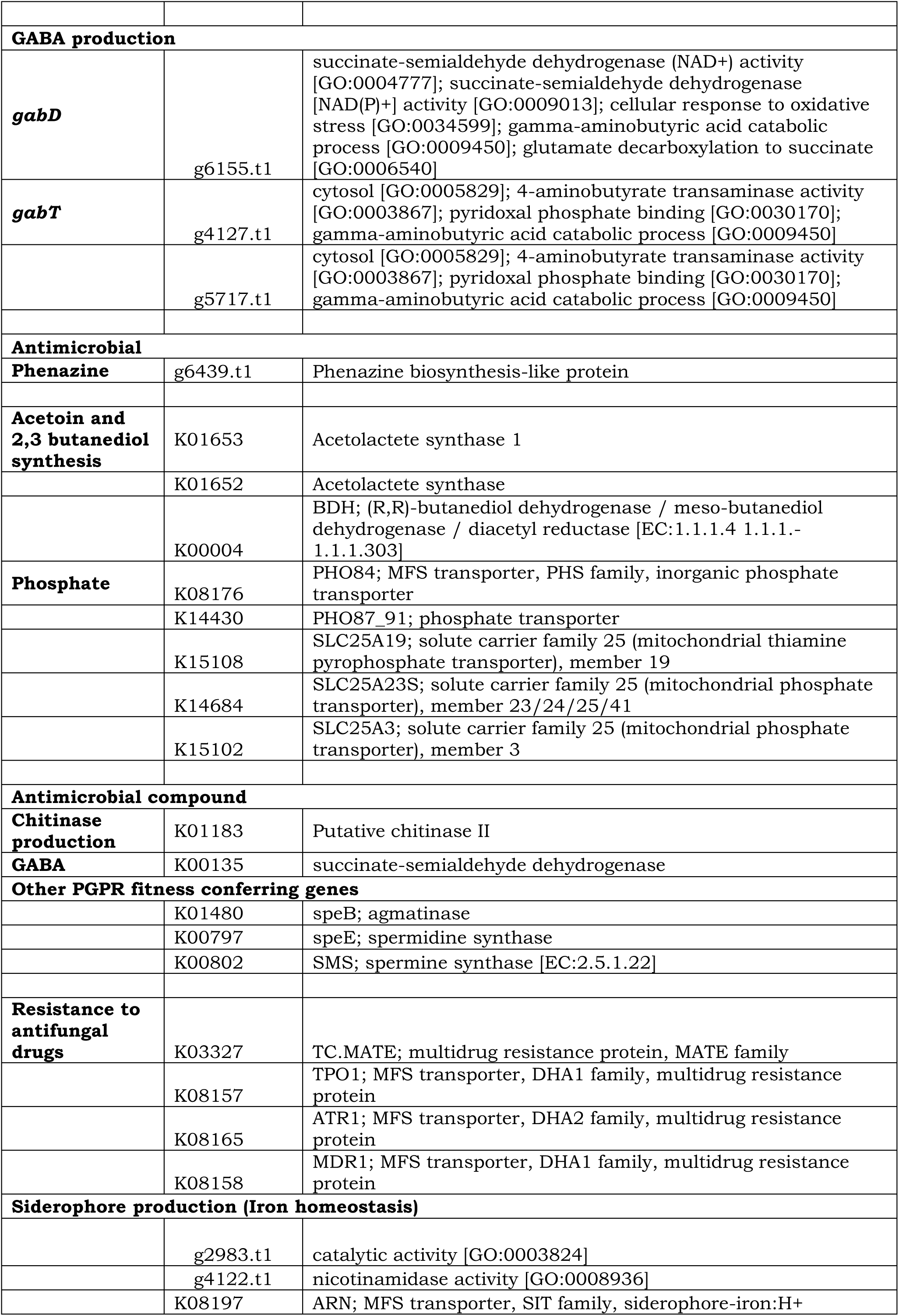
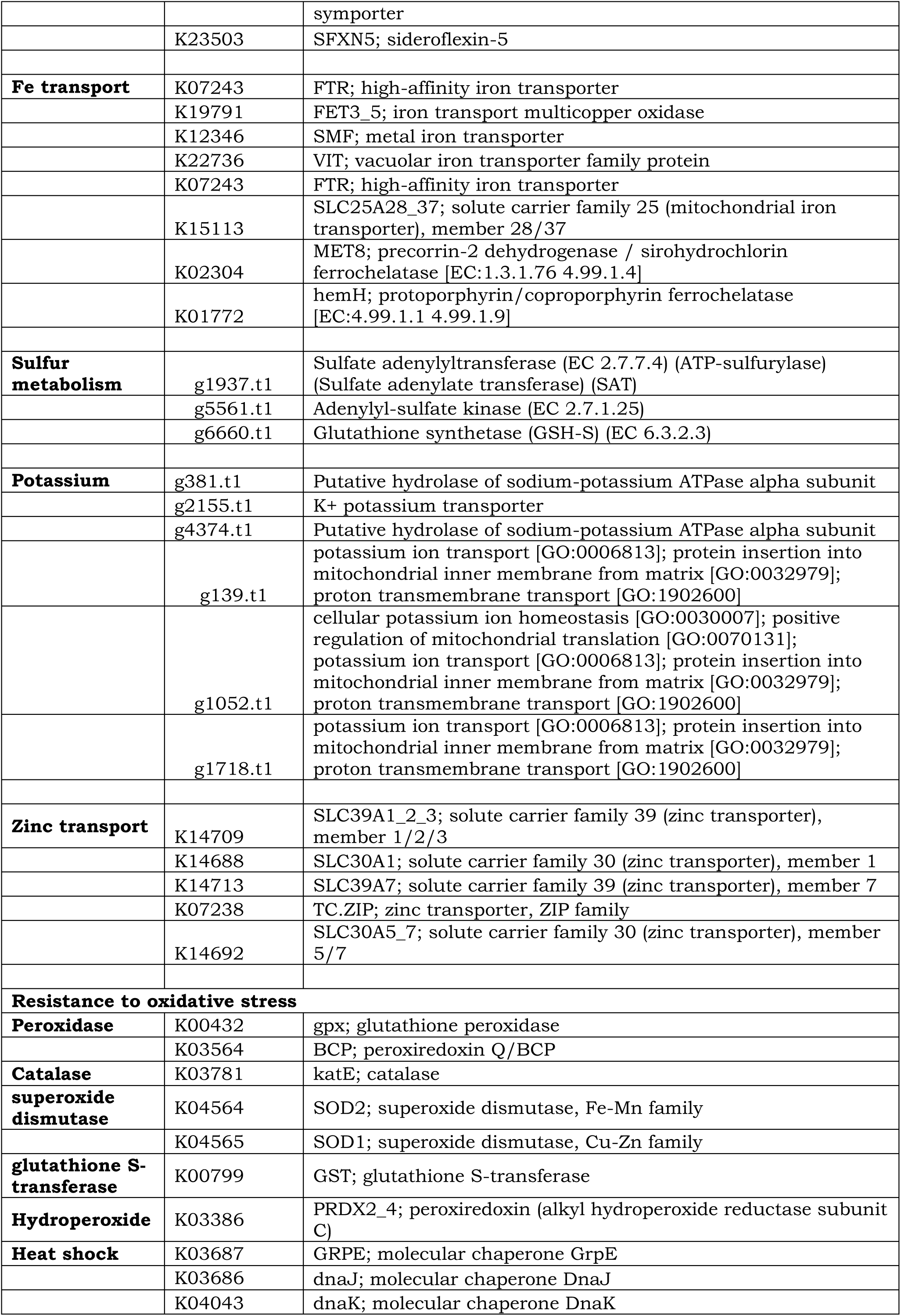
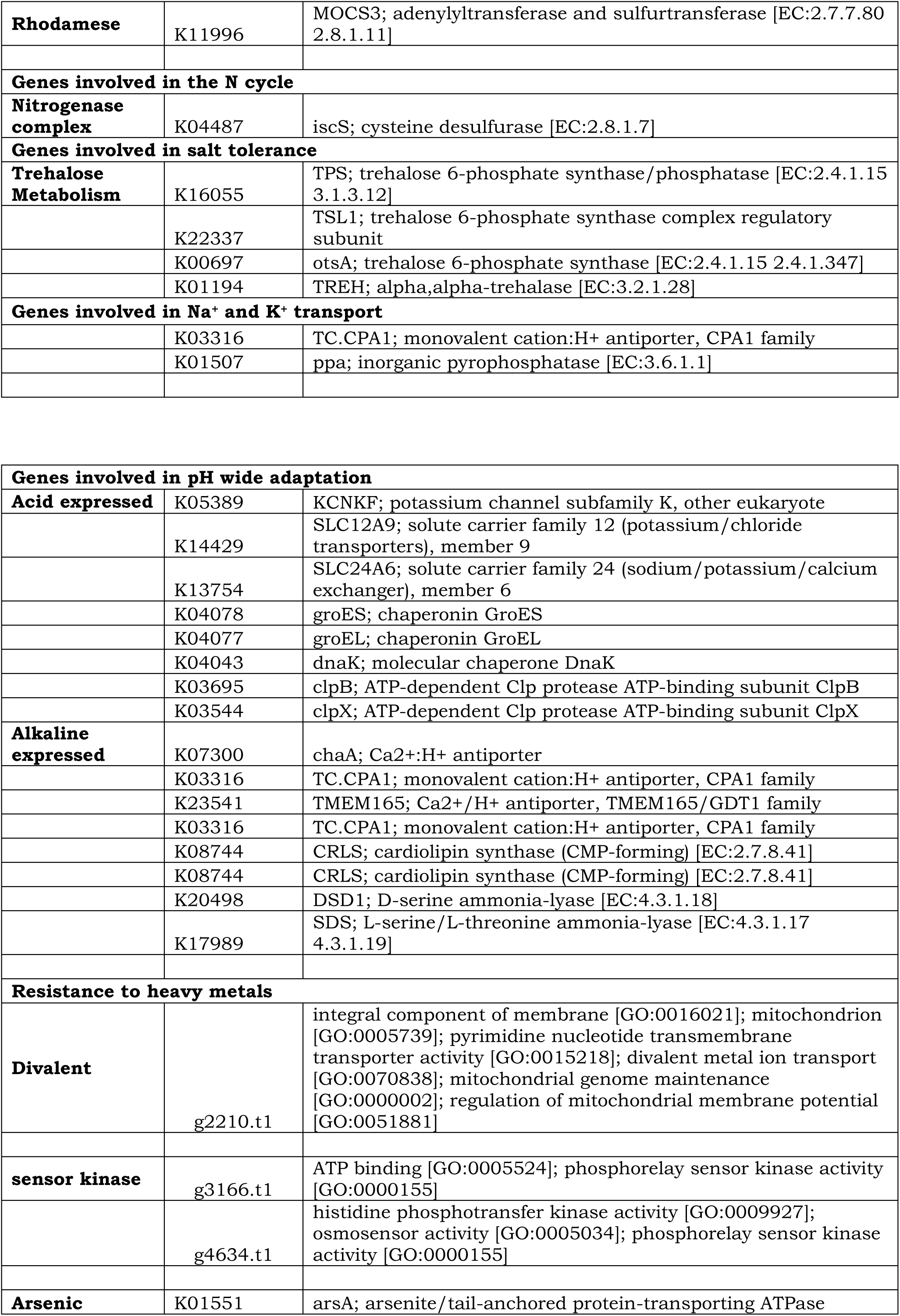
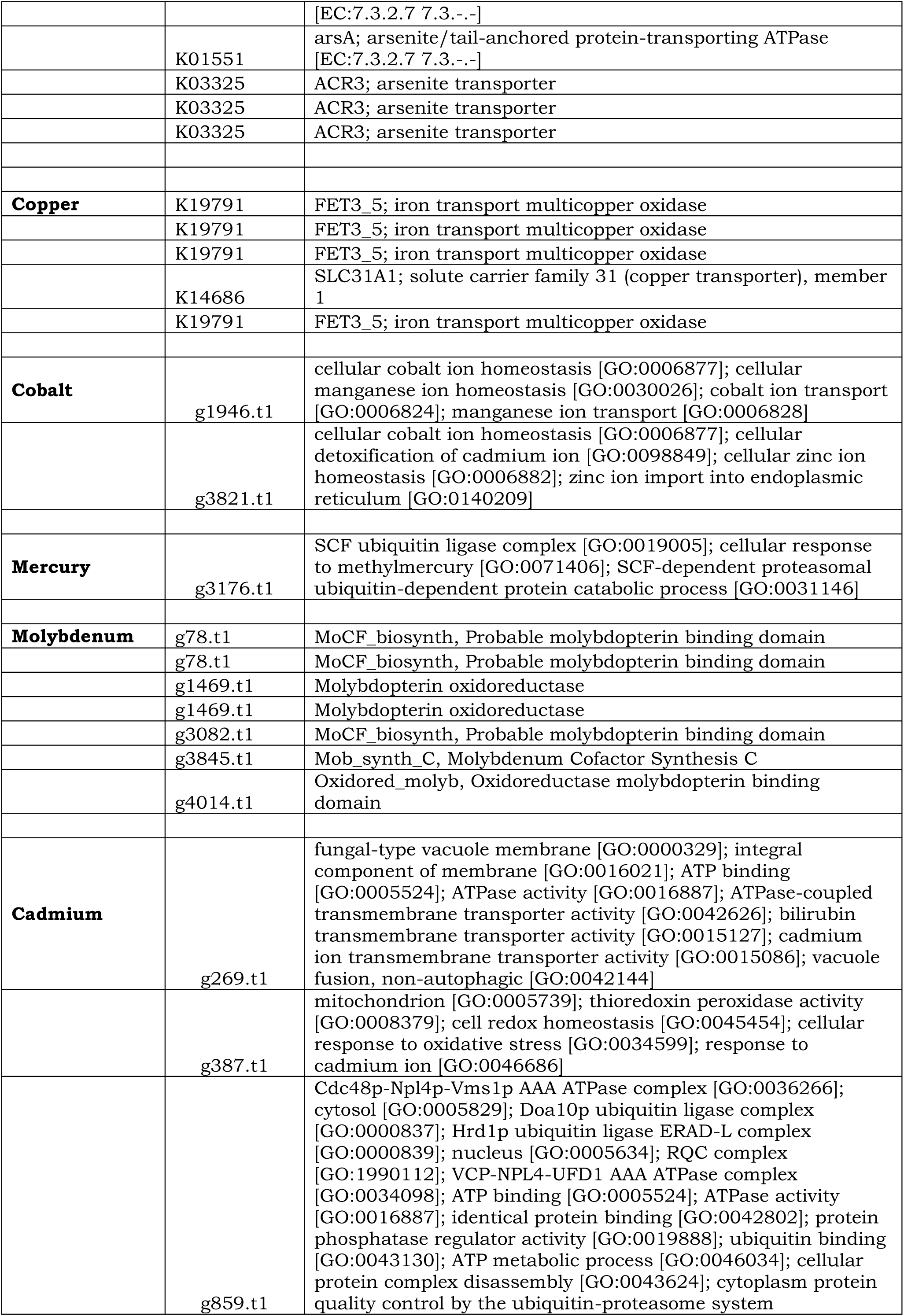
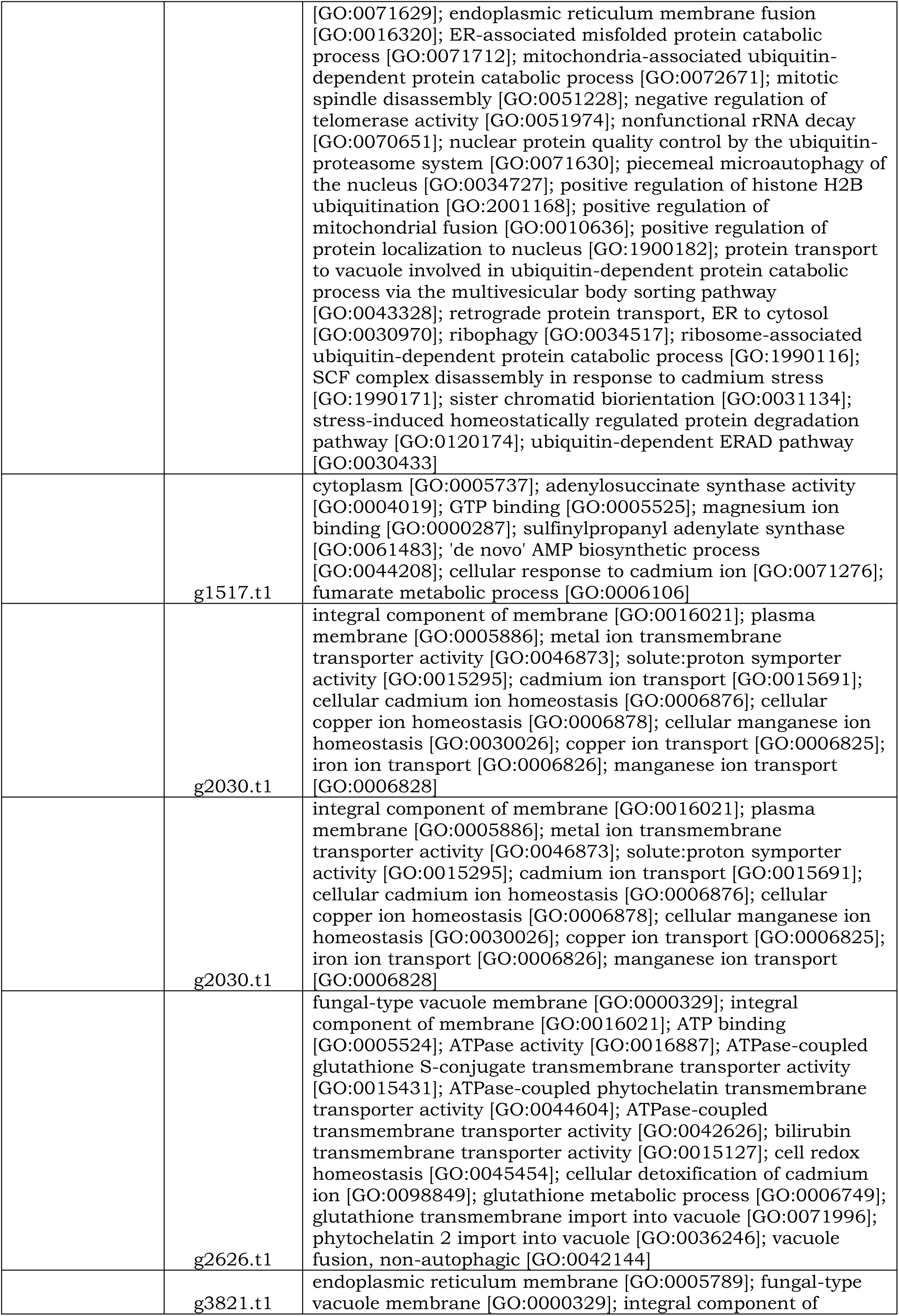
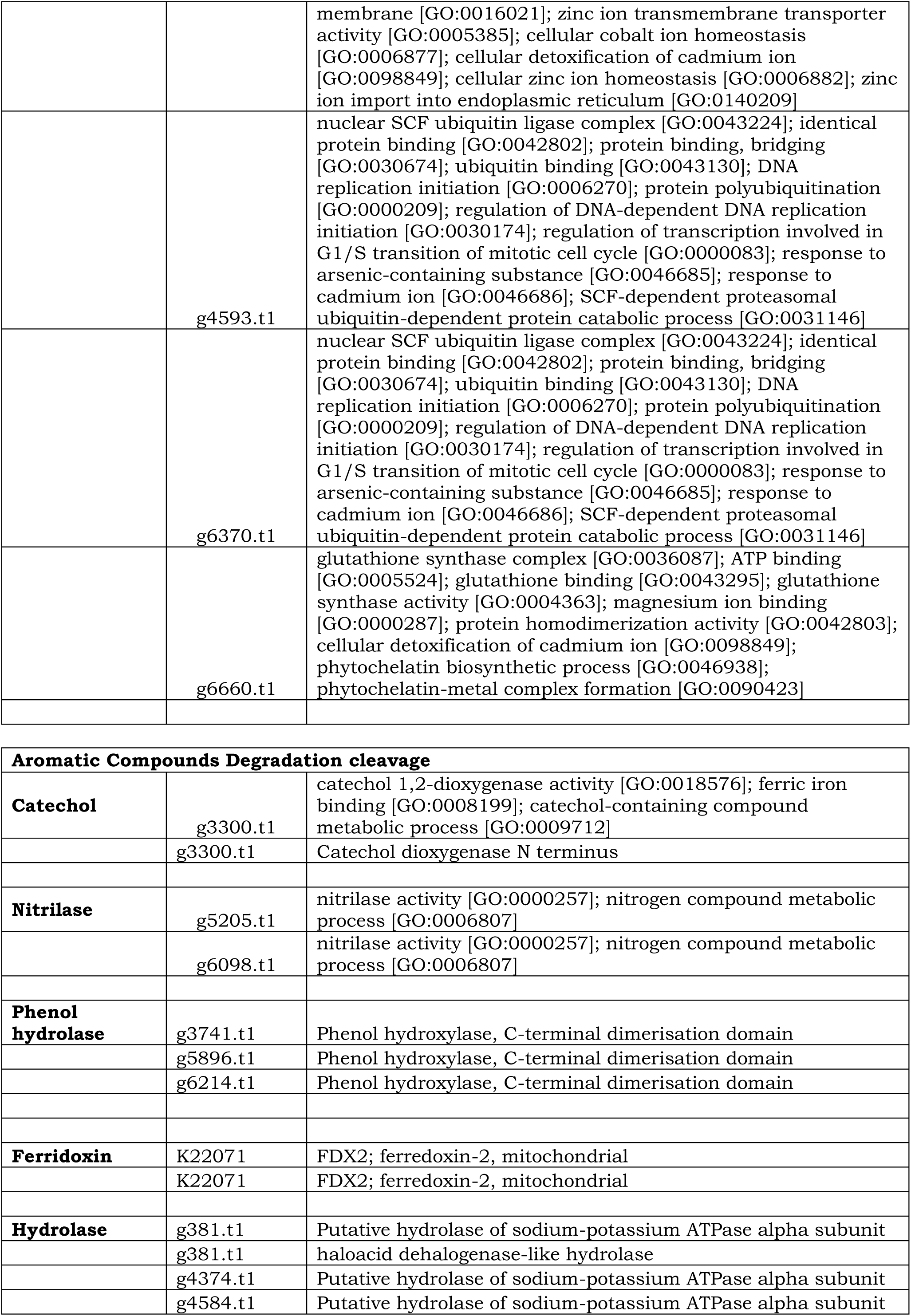
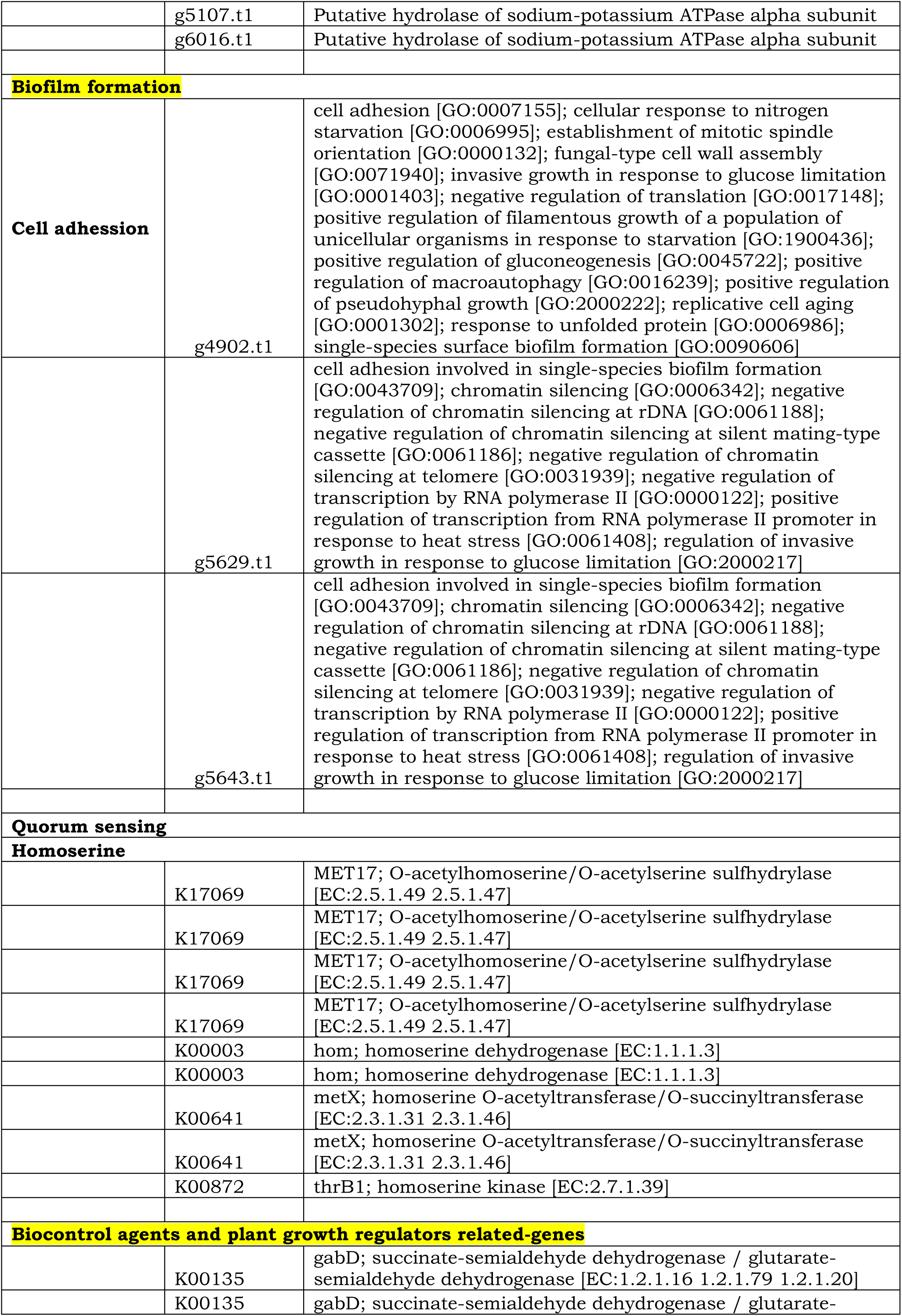
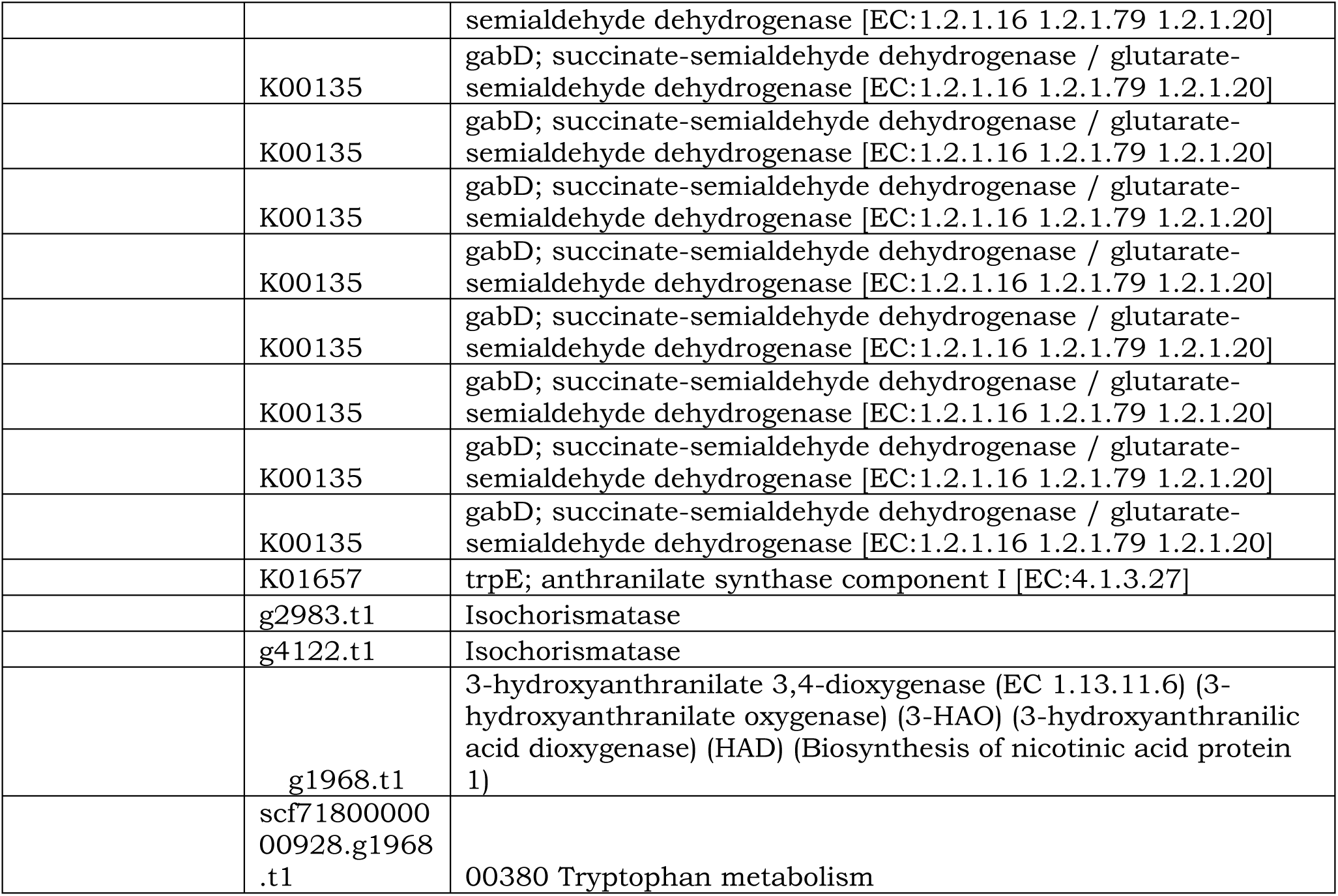
Genes related to Plant Growth Promoting traits that are annotated to be present in *Wickerhamomyces anomalus* strain MSD1

In addition to the above PGP traits, two growth-promoting **volatile organic compounds** (VOCs), acetoin and butanediol, were reported to promote plant growth by stimulating root formation and increasing systemic disease resistance and drought tolerance in some other very efficient PGPR[5,25–29]. Genes encoding enzymes including **acetolactate synthase** and **acetoin dehydrogenase** (Table 1) which are involved in acetoin and butanediol synthesis, were detected in the genome of MSD1 [24,28,30]. Two pyruvate molecules condensed into acetolactate is catalyzed by acetolactate synthase, and which is converted to acetoin by acetolactate decarboxylase and finally acetoin is converted to **2**,**3-butanediol** catalyzed by acetoin reductase [24,28,30].

### Nitrogen fixation

Nitrogenase is the enzyme central to nitrogen fixation and it consists of Fe-protein encoded by *nifH* and MoFe-protein encoded by *nifDK*. Full assembly of the nitrogenase complex needs the products of at least twelve nif genes, especially for the processing of catalytic stability and nitrogenase metalloclusters (nifMZ, nifUS, and *nifW*) and for synthesis of a particular molybdenum cofactor (MoCFC). Many microbial gene families are responsible for organic N decomposition, metabolism, and biosynthesis in soil. Here, five gene families directly related with N cycling processes were extracted and analyzed, including ***nao*** (nitroalkane oxidase), ***nmo*** (nitronate monooxygenase), gdh (glutamate dehydrogenase), *ureC* (urease) and **GS** (glutamine synthetase)[21].

MSD1 is able to grow on nitrogen-free medium (data not shown) and this indicates that the strain is able to fix atmospheric nitrogen. The MSD1 genome contains nif genes together with the *NifU* and nitronate genes which are the positive/negative regulatory proteins for *nif* genes [21].

### ACC Deaminase

One of the mechanisms of PGPR to alleviate salt stress is the synthesis of the enzyme 1-aminocyclo-propane-1-carboxylate (ACC) deaminase or its homologue D-cysteine desulfhydrase encoded by acdS or dcyD, respectively. Both enzymes lower ethylene accumulation in stressed plants by cleaving ACC, an immediate precursor of ethylene in plants, to form ammonia and α-ketobutyrate. This reaction is pyridoxal phosphate dependent, and both ACC deaminase and D-cysteine desulfhydrase belong to the pyridoxal phosphate-dependent enzyme family PALP. In the MSD1 genome, neither acdS genes nor dcyD genes are present but eight CDSs containing genes encoding genes belonging to the PALP domain (Table 1) was found [24,31,32]. Of these genes, presence of Cys_K 1 and 2 for Cysteine desulfurase (KO ID: K04487, 5 copies of iscS), Tryptophan synthase beta chain (2 copies) and L-threonine ammonia-lyase (6 copies) both show lyase activity and potentially perform ammonia synthesis similarly to the enzymes encoded by acdS and dcyD[30].

### Genes of Central Metabolism and Cellular Processes

Carbohydrate degradation pathways **Emden-Meyerhof pathway** and **Entner-Doudoroff pathway** for glucose, arabinose, mannose, trelose, mannitol, and the respective transport systems have been detected. All genes of the TCA cycle were present (**Table 1**). Exo- and polysaccharide biosynthesis and the respective transporter have been discovered (**Table 4**). A comprehensive list of detected genes is given in the **supplementary files**.

### Genes putatively involved in salt tolerance

MSD1 can grow well in 0–12% NaCl (data not shown). Analysis of the genome reveals that strain MSD1 has a number of genes related to salt tolerance. For example, trehalose can act as an osmoprotectant under environmental stresses such as high salt or drought, low temperature or osmotic stress in many organisms. Trehalose accumulates in transgenic rice and enhances plant abiotic stress tolerance. So far five trehalose biosynthetic pathways have been found in bacteria including treS, otsA/otsB, treP, treT and treY/treZ40[30,33]. Here, two trehalose biosynthesis pathway **otsA/B**, were identified in the MSD1 genome. In the otsA/otsB pathway both glucose-6-phosphate and UDP-glucose can synthesize trehalose-6-phosphate catalyzed by trehalose-6-phosphate synthase (otsA) activity. **Trehalose-6-phosphate** is then formed from trehalose catalyzed by trehalose-6-phosphate phosphatase (**otsB**) activity. Eventually, trehalose may be hydrolyzed by trehalase (2 copies) with the generation of two glucose molecules (**Table 4**). This pathway has been recognized as a universal pathway present in microorganisms and contributes to the survival under harsh environmental conditions [30,33].

Moreover, a number of **osmoregulation receptors** and transport systems were determined in the MSD1 genome. These genes can encode up to 24 two component systems (TCSs), among which 21 TCSs can be functionally assigned based on the KEGG database (**Table 4**) [30,33].

Of those 21 assigned TCSs, 3 belong to the SSK1 (response regulator) family, two to the YPD1 (phosphorelay intermediate protein) family, 16 belong to the SLN1 (sensor histidine kinase) family and one to the SKN7 (response regulator) family. The eight remaining TCS genes are annotated as sensor histidine kinase (**Table 4**).

In addition, genes encoding transport systems such as K+ transport systems for K+ accumulation and H+/Na+ antiporters (nha) for importing H+ and pumping out Na+ have also been found to resist hyperosmotic (Aft1 domain) stress in the genome of MSD1 (**Table 4**).

The MSD1 genome carries heat shock genes dnaJ, dnaK, groES, groEL, htpG, and grpE (**Table 4**). Moreover, the clpB gene, a **heat shock protein**, specified to be upregulated during salt stress in marine bacteria is also contained. The MSD1 genome also carries CDSs encoding for **peroxidases, superoxidase**, and glutathione S-transferase (**Table 1**). These genes play a role in the protection of cell **oxidative stress** caused by salt stress[30,33,34].

The genome sequence of marine isolate *W. anomalus* strain MSD1 presented in this paper is a plant growth promoting yeast isolated from the seaweed [12]. This study showed MSD1 has potential traits such as Zinc and phosphate-solubilizing, iron and sulfate transformation capability, production of ACC deaminase, siderophore, and VOCs; making it as an effective PGP yeast. Considering a variety of complex conditions that occur in rhizospheres [35], the environmental adaptability of PGPR in in situ rhizosphere became an important factor for improved plant growth-promoting capacity. In addition, initial studies focusing on the functional properties of PGPR have led to interest in the comparative analyses of pan-/core-genomes of these bacteria, which are of ecological importance for elucidating the fundamental genotypic features of PGPY [36,37].

## Conclusions

The genetic information obtained for *W. anomalus* strain MSD1 will enable us to interpret the expressed traits of the yeast and further provide insights into the practical applications of the strain as a bio-stimulant/PGPR for agriculture use or agri-input.

## Data availability

This whole genome sequence of the biosample SAMN12347843 has been deposited at GenBank/NCBI under the accession number SRR10092046 and BioProject number PRJNA556347. The associated Illumina HiSeq 4000 subreads are available under the SRA accession number SRR9822044. https://www.ncbi.nlm.nih.gov/bioproject/PRJNA556347.

## Acknowledgements

The work is an outcome of the collaborative BIRAC project between T. Stanes & Company Limited and DBT, India.

---

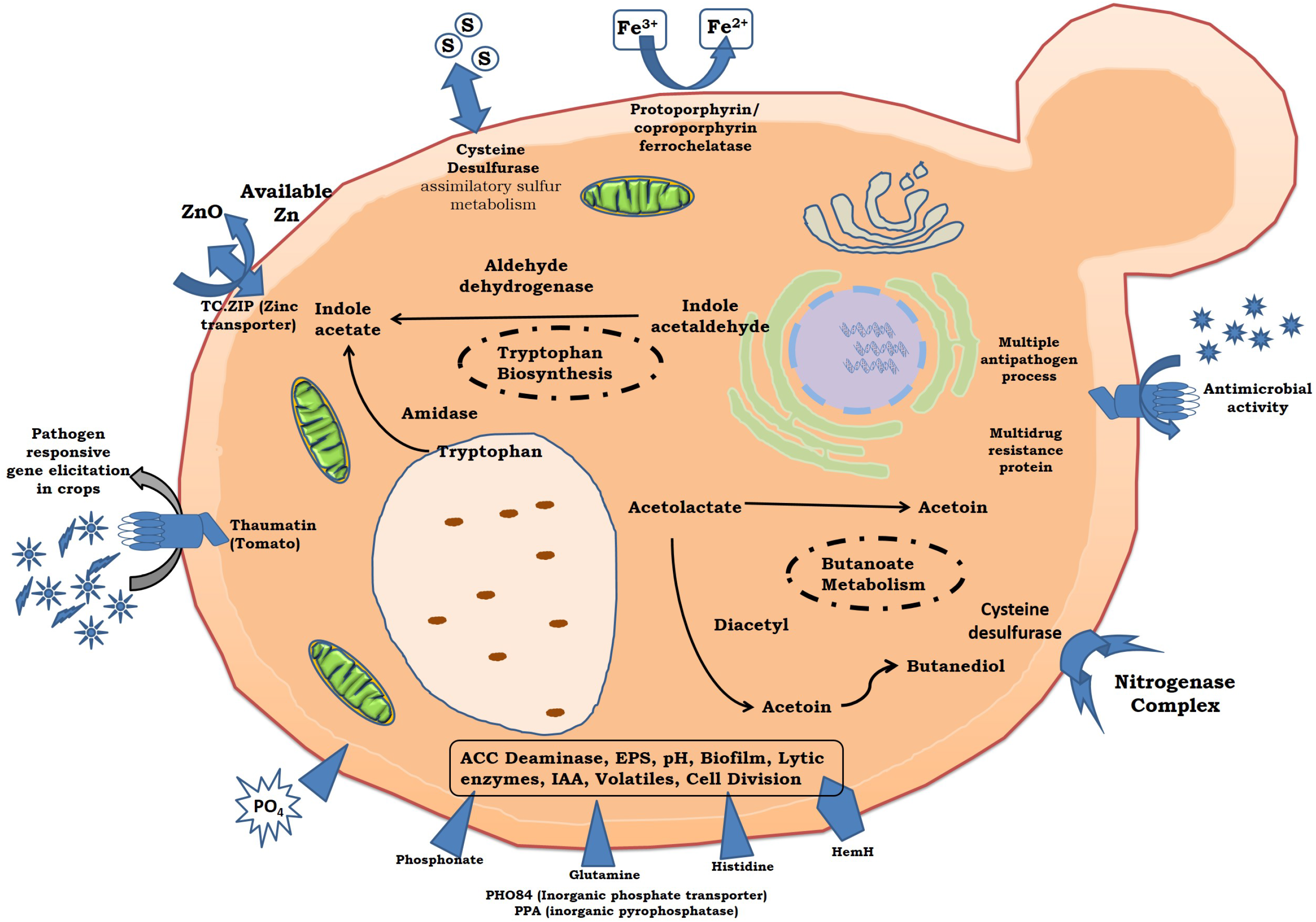

